# Timing of behavioral responding to long-duration Pavlovian fear conditioned cues

**DOI:** 10.1101/2023.01.25.525456

**Authors:** Kristina M. Wright, Claire E. Kantor, Mahsa Moaddab, Michael A. McDannald

**Affiliations:** Boston College, Department of Psychology & Neuroscience, Chestnut Hill, MA 02467

## Abstract

Behavioral responding is most beneficial when it reflects event timing. Compared to reward, there are fewer studies on timing of defensive responding. We gave female and male rats Pavlovian fear conditioning over a baseline of reward seeking. Two 100-s cues predicted foot shock at different time points. Rats acquired timing of behavioral responding to both cues. Suppression of reward seeking was minimal at cue onset and maximal before shock delivery. Rats also came to minimize suppression of reward seeking following cue offset. The results reveal timing as a mechanism to focus defensive responding to shock-imminent, cue periods.

## Introduction

Environmental events are organized by time and responding to these events is most advantageous when it takes event timing into account. Behavioral responding to associatively paired events not only reflects timing (Holland 2000; Jennings and Kirkpatrick 2006), but scalar expectancy theory proposes that timing is the basis of associative learning (Gallistel and Gibbon 2000; Kirkpatrick and Balsam 2016; Gibbon 1977). Behavioral demonstrations and brain investigations of timing primarily come from reward settings (MacDonald et al. 2013; Toso et al. 2021; Tallot and Doyère 2020; Buhusi and Meck 2005; Tam and Bonardi 2012), opposed to defensive settings (Mobbs et al. 2020). This is despite the fact that scalar expectancy theory is not specific to reward, and defensive responding is organized by time (Fanselow and Lester 1988).

The paucity of timing experiments in defensive settings, specifically Pavlovian fear conditioning, may stem from difficulty in demonstrating the behavioral phenomenon itself. For example, freezing to long-duration and short-duration Pavlovian fear conditioned cues shows little evidence of timing (Fanselow et al. 2019). Timing may be more readily observed when it must be balanced with other behaviors. In a conditioned suppression setting hungry rats responding for reward are intermittently played cues signaling foot shock. In this setting rats will suppress reward responding during cue presentation, but resume responding during non-cue periods (Estes and Skinner 1941). In support of timing, rats receiving unsignaled foot shocks at fixed intervals over a baseline of rewarded lever pressing will increase suppression of reward responding toward shock delivery (LaBarbera and Church 1974). Most pertinent, rats receiving Pavlovian fear conditioning to a long-duration cue over a baseline of rewarded lever pressing will acquire timing of defensive responding (Rosas and Alonso 1996). Suppression of rewarded lever pressing that is initially sustained over the entire cue duration becomes timed; minimal at cue onset, and increasing to cue offset (Rosas and Alonso 1996).

The current experiment had four goals. First, we set out to replicate Rosas and Alonso (1996) and demonstrate that timing of defensive responding emerges to long duration cues in a conditioned suppression setting. Second, we sought to determine if there are sex differences in timing. Third, we asked if timing of defensive responding would show a ‘peak-interval’ function (Buhusi and Meck 2006). That is, does increased defensive responding leading up to cue-signaled shock give way to decreased responding moving away from shock? Finally, we asked if timing was but one mechanism to focus defensive responding to cue periods when shock was imminent.

## Methods

Subjects were 16 adult Long Evans rats (8 females and 8 males) bred in the Boston College Animal Care Facility, and maintained on a 12-hr light cycle (lights off at 6:00 pm). Rats were individually housed with water freely available and specific food amounts provided daily. During behavioral testing, rats were restricted to 85% of their free-feeding body weight. All protocols were approved by the Boston College Animal Care and Use Committee and all experiments were carried out in accordance with NIH guidelines.

Behavioral testing took place in eight individual chambers with aluminum front and back walls, clear acrylic sides and top, and a grid floor. Chambers were contained within a sound attenuating shell. A nose poke above an external food cup were present at the center of one wall. Two speakers were mounted on the ceiling directly above the wall containing the food cup and nose poke. Rats were exposed to the experimental food pellets in their home cage, then shaped to nose poke for pellet delivery in the experimental chamber. Nose pokes were reinforced on a variable interval 60 s schedule, independent of Pavlovian contingencies.

The two 100-s, auditory cues to be used for Pavlovian fear conditioning consisted of repeating, 500 ms motifs of a horn or broad-band click, counterbalanced. Rats were exposed to each cue four times in a single session. During Pavlovian fear conditioning, each cue was paired with foot shock. The 62-min session consisted of four presentations of each cue (8 total presentations) with a mean inter-trial interval of 6 min. Cue order was randomly determined by the behavioral program. For the shock100 cue, foot shock (0.5 mA, 0.5 s) was delivered at cue offset, 100 s after cue onset (Figure 1A, top). For the shock50 cue, foot shock was delivered 50 s after cue onset (Figure 1A, bottom), and cue presentation continued for another 50 s.

**Figure 1.**
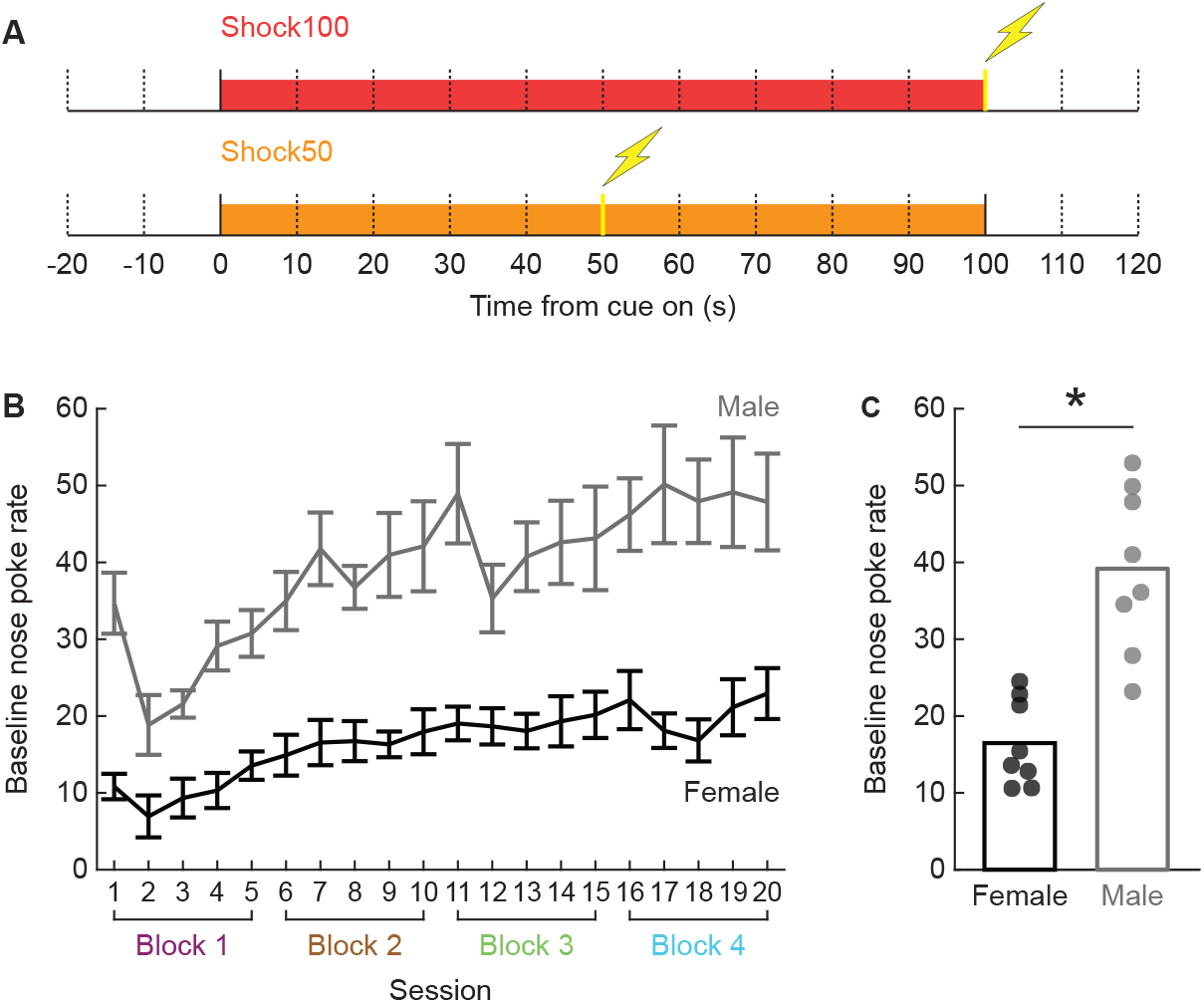
Experimental design and baseline nose poke rates. **(A)** Rats received 100-s cue presentations over a baseline of rewarded nose poking. Shock was delivered at the offset of the shock100 cue, while shock was delivered 50 s into the shock50 cue. Yellow vertical line and shock icon indicate time shock was delivered. **(B)** Mean ± SEM suppression ratio is shown for female (black) and male (grey) rats over the 20 Pavlovian fear conditioning sessions. Blocks are identified below the x-axis: 1 (magenta), 2, (brown), 3 (green), and 4 (cyan). **(C)** Group mean (bar) and individual (points) nose poke rate for all 20 sessions are shown. *independent samples t-test, p < 0.05.

Behavioral responding was measured with suppression ratio, calculated from nose poke rates taken in 10-s intervals starting 20 s prior to cue onset and continuing until 20 s following cue offset. The 10-s interval prior to cue onset served as baseline. Suppression ratio was calculated as: (baseline nose poke rate – interval nose poke rate) / (baseline nose poke rate + interval nose poke rate).

## Results

Male rats poked at higher baseline rates than female rats (Figure 1B). ANOVA [factors: sex (female and male) and session (1-20)] found significant main effects of sex and session, and a significant sex x session interaction (all F > 2, all *p* < 0.005). An independent samples t-test for mean nose poke rate across all sessions found a significant sex difference (t_14_ = 5.16, *p* = 0.00014; Figure 1C).

The 20 Pavlovian fear conditioning sessions were organized into four, 5-session blocks. Timing of behavioral responding to the shock100 and shock50 cues emerged over the four blocks (Figure 1C). During block 1 rats suppressed reward seeking continuously across cue presentation and after cue offset. Over blocks 2-4 rats reduced suppression of reward seeking during cue onset, maintained suppression just before shock delivery, then decreased suppression after cue offset. No sex differences were observed. In support, ANOVA for mean suppression ratio [factors: sex (female and male), cue (shock100 and shock50), block (1-4), and time (14, 10-s intervals from 20 s prior to 20 s following cue offset)] found a significant block x time interaction (F_39,546_ = 5.34, *p* = 2.21 × 10^−20^), and a significant cue x block x time interaction (F_39,546_ = 3.40, *p* = 1.48 × 10^−10^). No significant main effect of sex, or significant sex interactions were observed (all F < 1.3, all *p* > 0.15).

To reveal cue-specific patterns we performed separate ANOVAs for the shock100 and shock50 cues, removing sex as a factor. ANOVA for shock100 [factors: block (1-4) and time (10, 10-s cue intervals)] found a significant block x time interaction (F_27,405_ = 5.11, *p* = 4.29 × 10^−14^). The interaction was driven by reduced suppression ratios to the first 10-s cue interval during blocks 2-4 (Figure 2A). Compared to block 1, 95% bootstrap confidence intervals found lower suppression ratios during the first 10-s cue interval for block 2 (95% bootstrap confidence interval lower bound (LB) = 0.09, upper bound (UB) = 0.42), block 3 (LB = 0.27, UB = 0.58), and block 4 (LB = 0.41, UB = 0.69; Figure 2B, left). By contrast, suppression ratios for the 10-s interval prior to shock onset did not differ across blocks (all 95% bootstrap confidence intervals contain zero; Figure 2B, middle). Mirroring the pattern in the first 10-s interval, immediate post cue suppression ratios declined during blocks 2, 3, and 4 compared to block 1 (all 95% bootstrap confidence intervals did not contain zero; Figure 2B, right).

**Figure 2.**
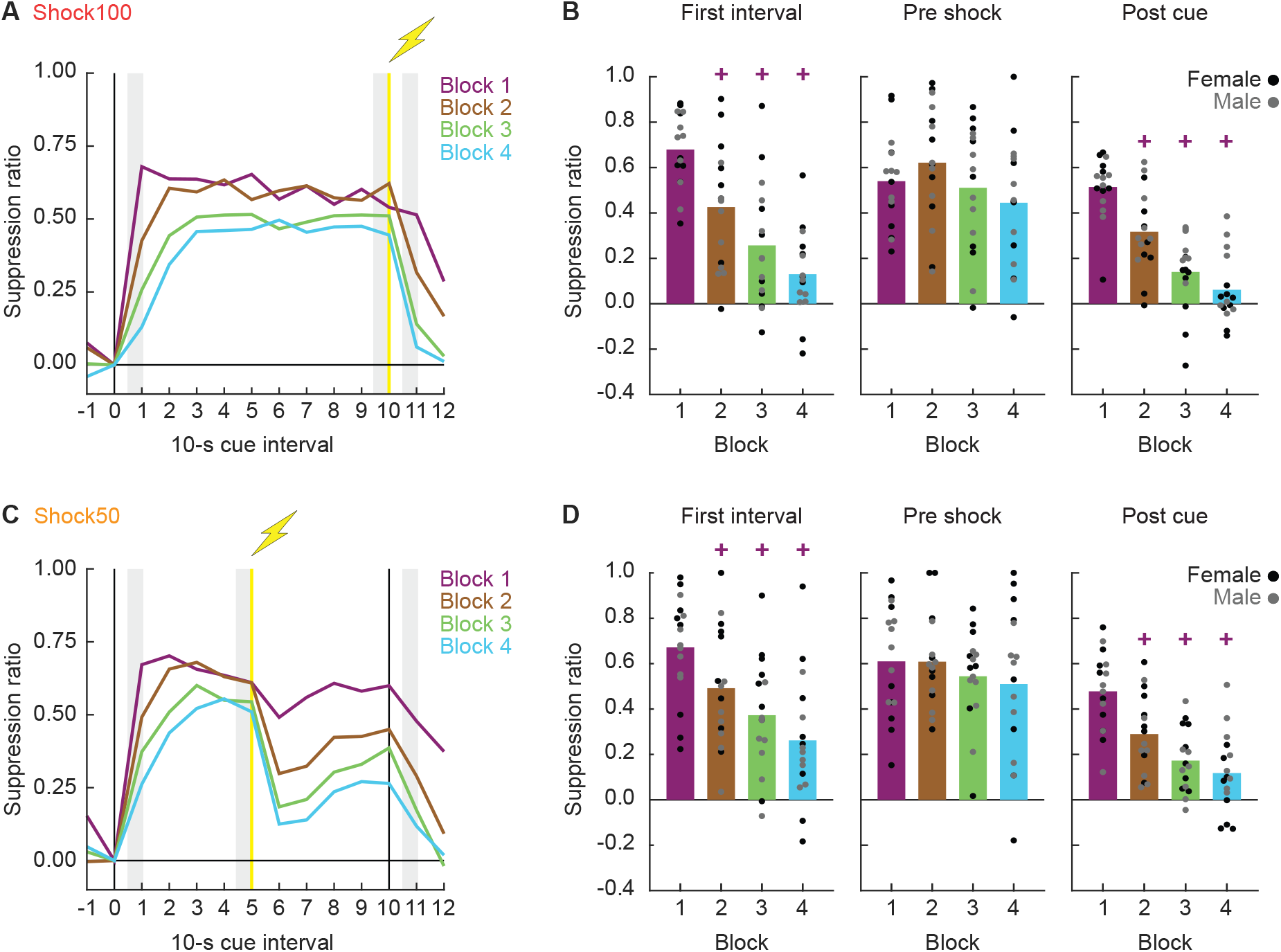
Timing of responding to Pavlovian fear conditioned cues. **(A)** Mean suppression ratio is shown for the 14, 10-s intervals around shock100 cue presentation for blocks 1 (magenta), 2, (brown), 3 (green), and 4 (cyan). Yellow vertical line and shock icon indicates time shock was delivered. **(B)** Group (bar) and individual (points) mean suppression ratios are shown for the first cue interval (left), pre shock interval (middle), and first post cue interval (right) during blocks 1-4. Females are black points and males are grey points. The three shaded regions in A correspond to the three trial periods in B. **(C)** Mean suppression ratio around shock50 presentation shown as in A. **(D)** Group (bar) and individual (points) mean suppression ratios shown as in B. +95% bootstrap confidence interval does not contain zero, each block is compared to block 1.

The temporal pattern of responding to the shock50 cue reinforces the narrative that over the four blocks, suppression of reward seeking became more specific to cue periods during which shock was imminent. ANOVA for the first 50 s of the shock50 cue [factors: block (1-4) and time (first 5, 10-s cue intervals)] found a significant block x time interaction (F_12,180_ = 4.82, *p* = 7.66 × 10^−7^). The interaction was driven by reduced suppression ratios to the first 10-s cue interval during blocks 2-4 (Figure 2C). 95% bootstrap confidence intervals found lower suppression ratios during the first 10-s cue interval for block 2 (LB = 0.05, UB = 0.30), block 3 (LB = 0.15, UB = 0.45), and block 4 (LB = 0.26, UB = 0.56), compared to block 1 (Figure 2D, left). Suppression ratios for the 10-s interval prior to shock onset did not differ across blocks (all 95% bootstrap confidence intervals contain zero; Figure 2D, middle). Post cue suppression ratios declined during blocks 2, 3, and 4 compared to block 1 (all 95% bootstrap confidence intervals did not contain zero; Figure 2D, right).

The results so far replicate and extend the findings from Rosas et al. 1996. Timing of behavioral responding (nose poke suppression) to long-duration Pavlovian fear conditioned cues is evident in female and male rats. Reduced responding at both cue onset and immediately following cue offset indicates an overarching trend to confine suppression of reward seeking to cue periods in which shock is imminent. Next we asked if we observed the second half of the peak-interval effect – reduced responding moving away from shock presentation on shock50 trials. We focused on suppression ratios from the last 50 s of the shock50 cue (Figure 3A), during which the cue continued to play after the shock had been delivered.

**Figure 3.**
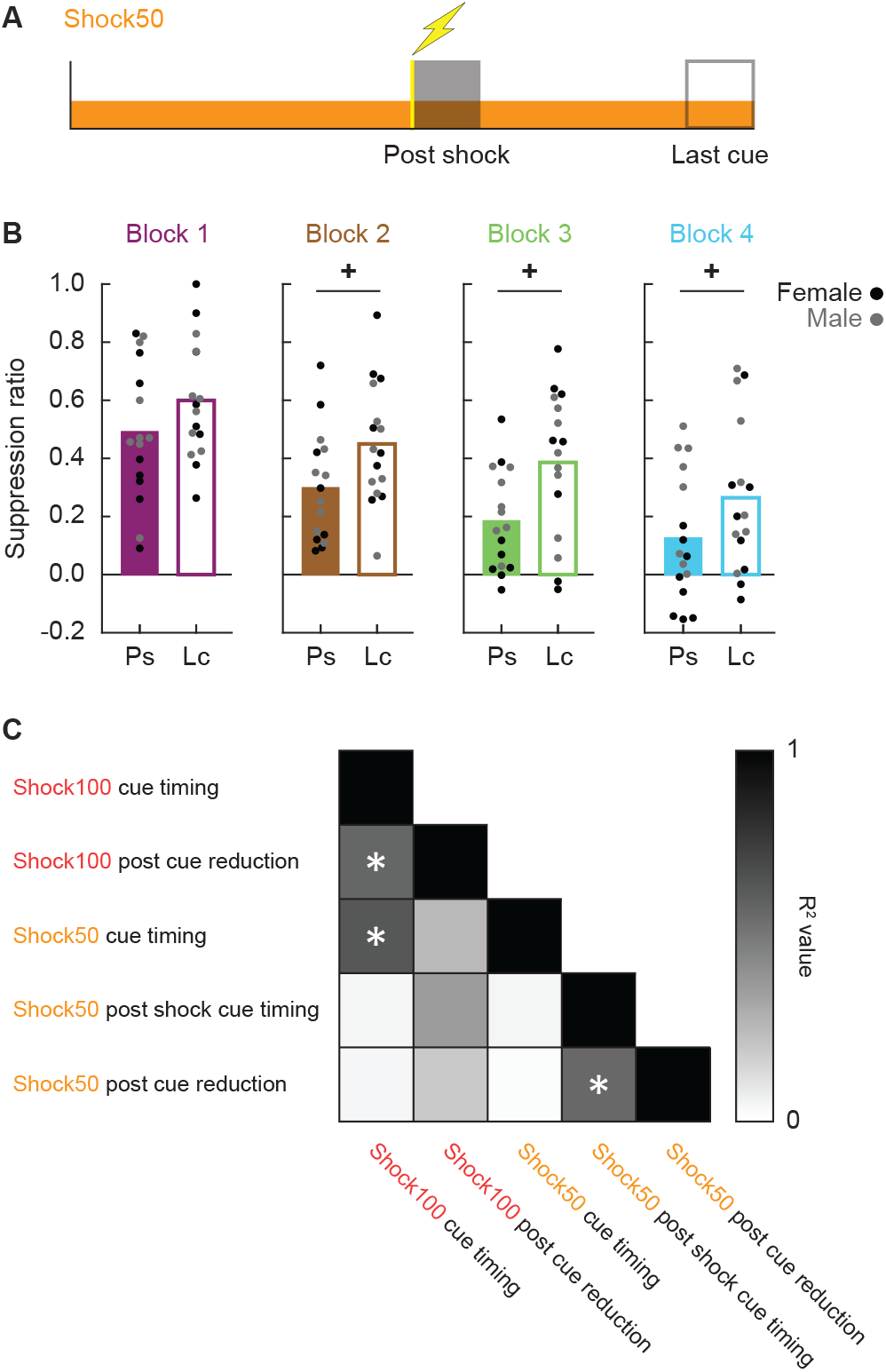
Timing of responding following shock suppression of reward and correlated temporal responding. **(A)** The diagram shows the post shock interval (closed bar) and last cue interval (open bar) for the shock50 cue. Yellow vertical line and shock icon indicate time shock was delivered. Females are black points and males are grey points. **(B)** Group (bar) and individual (points) mean suppression ratios are shown for the post shock cue interval (Ps, closed bar) and the last cue interval (Lc, open bar) during blocks 1 (magenta), 2, (brown), 3 (green), and 4 (cyan). **(C)** A correlation matrix was constructed for the five instances of temporal responding. Pearson’s correlation coefficient (R2) for each comparison is visualized from 0 (white) to 1 (black). *R^2^ *p*-value < 0.005 (Bonferroni correction for 10 comparisons).

Against our hypothesis, ANOVA [factors: block (1-4) and time (last 5, 10-s cue intervals)] found a significant main effect of block (F_3,45_ = 15.1, *p* = 6.15 × 10^−7^) a significant main effect of time (F_4,60_ = 7.12, *p* = 9.3 × 10^−5^), but no significant block x time interaction (F_12,180_ = 0.69, *p* = 0.76). The main effect of block was driven by an overall reduction in suppression from block 1 to block 4 (Figure 3B, left to right). However, the main effect of time was driven by *increased* suppression from the post shock cue interval to the last cue interval (Figure 3B, closed vs. open bars). 95% bootstrap confidence intervals revealed greater suppression during the last cue period compared to the post shock cue period for block 2 (LB = 0.04, UB = 0.27), block 3 (LB = 0.07, UB = 0.33), and block 4 (LB = 0.03, UB = 0.25). The results reveal a second instance of timing rather than the second half of a peak-interval effect: increasing suppression of reward seeking away from shock delivery and towards cue offset.

By block 4 we detected five, separate changes in temporal responding around shock100 and shock50 cue presentation. For both cues we observed timing of responding from cue onset to shock delivery, and reduced responding following cue offset. Timing of post-shock cue responding was additionally observed during shock50 presentation. To determine if changes in temporal responding were related, we calculated individual difference scores from the relevant time periods (shaded regions of Figure 2A, C) and constructed a correlation matrix (Figure 3C). Individuals showing greater timing of responding over shock100 presentation showed greater timing over shock50 presentation (R^2^ = 0.57, *p* = 2.47 × 10^−4^), as well as greater reductions in responding following shock100 offset (R^2^ = 0.63, *p* = 7.34 × 10^−4^). Separately, individuals showing greater timing of responding over the last half of shock50 cue presentation (as in Figure 3B, far right) showed greater reductions in responding following shock50 cue offset (R^2^ = 0.56, *p* = 8.28 × 10^−4^). Timing of responding over the last half of shock50 cue presentation was unrelated to initial timing of responding for shock50 (R^2^ = 0.05, *p* = 0.42) and shock100 (R^2^ = 0.05, *p* = 0.41).

## Discussion

The results provide conclusive answers to our first two questions concerning replication and biological sex. The emergence of timing to the shock100 cue replicates work by Rosas and Alonso (1996). Equivalent timing was observed in females and males, supporting timing of defensive responding as a core behavioral mechanism in adult rats. These results are consistent with findings that cue onset periods distant from aversive outcome can acquire inhibitory associative properties, while cue periods adjacent to shock outcome can acquire excitatory properties (Rescorla 1967, 1968).

We did not observe peak-interval responding, which would have been a reduction in responding moving away from shock presentation (on shock50 trials). Instead we observed an increase in responding during the 50-s, post-shock cue period. This increase did not purely reflect generalization of timing during the initial 50-s cue period, as there was no relationship between changes in responding across individuals. The full pattern of post-shock cue responding may reflect two opposing mechanisms. A fear reduction mechanism may normally work to reduce responding following shock offset, evidenced by the overall decrease in responding across blocks. A working memory mechanism may be required to accurately track the current cue period: pre shock vs. post shock. Difficulty in cue period tracking would be expected to increase over the 50-s, post-shock cue presentation; pitting post-shock mechanisms to reduce responding against pre-shock mechanisms to increase responding over cue presentation.

If our results are so robust, why isn’t timing of defensive responding more commonly reported? We think the answer is equal parts procedure and adaptive behavior. We assume the goal of defensive systems is survival. In a traditional Pavlovian fear conditioning procedure, rats receive a small number of cue-shock pairings (typically auditory cues and long/intense foot shocks) in a neutral chamber, then testing responding in a novel chamber. Austere chamber design means there is little to do in these contexts. Rats then have limited information about their contexts and few behavioral options inside them. When a shock-predictive cue is played in these contexts, behavioral nuance such as timing of responding is not adaptive.

We used shorter and less intense foot shocks that can produce lesser freezing and permit additional behaviors to be observed (Holland 1979; Mitchell et al. 2022; Chu et al. 2022; Gruene et al. 2015). Further, our rats had extensive, reward experience in the context. Like Fanselow and colleagues (2019) we found that initial responding to fear conditioned cues showed no evidence of timing. However, continuing this pattern would mean spending large periods of time hungry, but not poking for food pellets. In a conditioned suppression setting, adaptive responding means confining defensive responding to periods when the aversive outcome is imminent. Consistent with this description, we found that timing was one of several, related mechanisms to focus defensive responding to shock-imminent cue periods.

Using a procedure in which distinct cues predict unique foot shock probabilities (0, 0.25 or 1) we have found that generalized cue responding gives way to responding that scales to each cue’s shock probability (Ray et al. 2020; Walker et al. 2018; Moaddab et al. 2021). This pattern, and the timing pattern we observed here, has parallels to stress and anxiety disorders. One way to conceptualize disordered fear and anxiety is that adaptive defensive responding becomes maladaptive, by being displayed at inappropriate times or to non-threatening stimuli. Understanding how generalized defensive responding transitions to specific responding that utilizes timing and threat probability may provide insight into the behavioral and neural basis of stress and anxiety disorders.

## Acknowledgements

Research reported in this publication was supported by the National Institute of Mental Health of the National Institutes of Health under Award Number MH117791. The content is solely the responsibility of the authors and does not necessarily represent the official views of the National Institutes of Health. The authors report no competing interests.

